# PGSbuilder: An end-to-end platform for human genome association analysis and polygenic risk score predictions

**DOI:** 10.1101/2023.04.12.536584

**Authors:** Ko-Han Lee, Yi-Lun Lee, Tsung-Ting Hsieh, Yu-Chuan Chang, Su-Shia Wang, Geng-Zhi Fann, Wei-Che Lin, Hung-Ching Chang, Ting-Fu Chen, Peng-Husan Li, Ya-Ling Kuo, Pei-Lung Chen, Hsueh-Fen Juan, Huai-Kuang Tsai, Chien-Yu Chen, Jia-Hsin Huang

## Abstract

Understanding the genetic basis of human complex diseases is increasingly important in the development of precision medicine. Over the last decade, genome-wide association studies (GWAS) have become a key technique for detecting associations between common diseases and single nucleotide polymorphisms (SNPs) present in a cohort of individuals. Alternatively, the polygenic risk score (PRS), which often applies results from GWAS summary statistics, is calculated for the estimation of genetic propensity to a trait at the individual level. Despite many GWAS and PRS tools being available to analyze a large volume of genotype data, most clinicians and medical researchers are often not familiar with the bioinformatics tools and lack access to a high-performance computing cluster resource. To fill this gap, we provide a publicly available web server, PGSbuilder, for the GWAS and PRS analysis of human genomes with variant annotations. The user-friendly and intuitive PGSbuilder web server is developed to facilitate the discovery of the genetic variants associated with complex traits and diseases for medical professionals with limited computational skills. For GWAS analysis, PGSbuilder provides the most renowned analysis tool PLINK 2.0 package. For PRS, PGSbuilder provides six different PRS methods including Clumping and Thresholding, Lassosum, LDPred2, GenEpi, PRS-CS, and PRSice2. Furthermore, PGSbuilder provides an intuitive user interface to examine the annotated functional effects of variants from known biomedical databases and relevant literature using advanced natural language processing approaches. In conclusion, PGSbuilder offers a reliable platform to aid researchers in advancing the public perception of genomic risk and precision medicine for human disease genetics. PGSbuilder is freely accessible at http://pgsb.tw23.org.

## Introduction

An ultimate goal of human genetics is to understand the genetic basis of human diseases, diagnosis, and management. Results from a large amount of genome-wide association studies (GWAS) have vastly demonstrated that many single nucleotide polymorphisms (SNP) genetic variants are associated with various complex traits^1^. In early 2023, more than 6,300 studies have conducted to map over 496,000 associations between human SNPs and diseases/traits in the GWAS catalog^2^. In the past two decades, the successes of GWAS not only drive the discovery of deleterious mutations linked to certain disease phenotypes but also imply a general pattern of polygenicity of common diseases^3, 4^. Many common diseases that conform to polygenic inheritance are underpinned by multiple genetic variants with small or moderate effects^5^. After the realization of a large proportion of the variance in genetic liability to common diseases, utilization of causative risk alleles based on the GWAS discoveries for disease risk prediction has become the potential to stratify patients for precision prevention^6, 7^.

Polygenic risk score (also known as polygenic scores; PRS) is an important methodology to leverage the genetic contribution of an individual’s genotype to measure the genetic liability to complex traits or diseases^8, 9^. Clumping and thresholding (C+T)^10^ is the primary PRS method based on the summary statistics from GWAS by pruning SNPs through a process of Linkage Disequilibrium (LD) clumping and selecting a *P*-value threshold. Still, it has limitations in the predictive performance without considering other genetic factors. Currently, several PRS methods based on the summary statistics apply a different selection of the prior distribution on the effect sizes of the SNPs under the Bayesian framework. For example, LDpred^11^ and LDpred2^12^ improve the prediction performance by enhancing LD modeling based on the normality assumption. PRS-CS^13^ introduces a different concept to provide a continuous shrinkage (CS) prior to accommodate diverse underlying genetic architectures. Alternatively, SBayesR^14^ and SDPR^15^ assume a different mixture of normal distributions on the individual-level data as input for adaptive modeling of SNP effect size. Lassosum^16^ implements a penalized regression approach with a Lasso-type penalty. Empirical evidence from benchmark experiments shows that not a single method clearly outperforms all other methods in the prediction accuracy for all the simulated data and disease traits^12–14, 17^. Nevertheless, each different PRS method can potentially improve the development of PRS construction with specific optimization procedures. Recent studies have demonstrated that the comparison of many PRS methods could facilitate the future implementation of PRS in clinical settings^18, 19^. Although a few practical guidelines have introduced how best to perform PRS analyses^20–22^, a steep learning curve of implementing those PRS packages and the computing resources required by some tools are impractical for doctors and clinical professionals.

As the popularity of PRS increases, over 400 publications report more than 3,200 polygenic scores in the Polygenic Score Catalog (https://www.PGSCatalog.org)^23^. However, those PRS studies were predominantly conducted on individuals of European descent^24^. Due to the poor transferability of PRS across populations^25, 26^, one critical step toward effectiveness in PRS accuracy is to conduct PRS development for the diversity of participants from different ancestries. Along with the cost of a single genetic test per individual plummeting to less than US$50, it becomes feasible to acquire a sufficient cohort size for PRS from the population with underrepresented ancestries by the medical institutes in different countries. In addition, the current consensus about the refinement of PRS should include other informative clinical factors based on their healthy records. To facilitate genetic analysis and PRS development, a sophisticated analysis platform could enable the construction of PRS in clinical research efficiently. For example, impute.me is a recently developed web tool to provide basic PRS estimation using a single method of LDpred to predict individual polygenic risks^27^. To increase the clinical practice of PRS, a comprehensive comparison of different PRS methods could leverage the extent of predictive values into a better understanding of the genetic liability for disease traits.

In this study, we present PGSbuilder which is an integrated cloud-based platform to analyze human genotype data. PGSbuilder provides a one-stop service to conduct both GWAS and PRS analyses and interactively visualize the analysis results. In PGSbuilder, users can run six different PRS methods as well as the PRS models with clinical factors to compare their performances concurrently. To the best of our knowledge, no other existing web server offers the possibility to compare multiple PRS models. Further, the interpretation of PRS is needed to apply the scores into biological explanations and clinical use. Notably, PGSbuilder also integrates the variant annotation automatically for the candidate SNPs from GWAS and PRS analyses using Ensembl Variant Effect Predictor (VEP)^28^ and biomedical literature mining from pubmedKB^29^. In addition, our web interface allows easy access to link all genetic analysis results and candidate SNP information with interactive displays. Finally, users can download all the analysis output files for further exploration.

## Materials and Methods

### Data privacy and security

Because genetic data will be uploaded to our server, a wide array of security measures are in force to ensure data privacy and security. Our local server has ISO 27001 certification for implementing an information security management system (ISMS). In addition, our server is designed based on the express MVC (Model-View-Controller) framework that encapsulates our features surrounded by powerful security layers. All interactions with the server are protected and secured with HTTPS. Any input data is deleted from our server once the analysis is completed. With the encryption by a firm one-time password, all analyzed results can only be accessed by the data uploader via an encrypted connection, within a 14 days timeframe.

### GWAS

To conduct quality control (QC) procedures and following genome-wide association studies (GWAS), we utilize PLINK 2.0, a comprehensive genome association analysis tool for population genetics^30^. There are three major steps for QC and two for GWAS. QC consists of variant filtering, individual filtering, and population stratification while GWAS analysis consists of principal component analysis (PCA) and association test.

First, unqualified SNPs are filtered out according to the minor allele frequency, Hardy-Weinberg equilibrium, and missingness. Secondly, individuals with the high missing rate of SNPs, large deviation of heterozygosity rate, and high kinship coefficient^31^ are also removed. Finally, to exclude individuals with different populations, population stratification is conducted against the population in HapMap 3^32^. Most of the QC criteria and recommended thresholds are referred to Marees et al^33^.

For the GWAS analysis, the top 10 principal components extracted from PCA are used to correct the genetic difference between in-group individuals^34^. Of note, the population stratification during the QC analysis is also conducted via PCA to remove outliers at the level of population, such as Asians, Africans, or Europeans. Next, the principal components and other provided covariates are included to correct the genetic effect during association tests. Only the effect size of autosomal SNPs is calculated using the “glm” function of PLINK 2.0^30, 35^.

### PRS methods

In PGSbuilder, the input dataset is separated into the base, target, and test sets, respectively. First, QC is applied on both base and target sets, and then GWAS is only performed on the base set to get the summary statistics. Combining the summary statistics with the target set which is used for the calculation of linkage disequilibrium (LD) and the selection of hyperparameters, PGSbuilder performs PRS analysis to build models based on different methods. This pipeline of PRS analysis is referred to Choi et al^21^. There are six PRS methods provided in PGSbuilder, including clumping and thresholding, PRSice2, LDpred2, Lassosum, PRS-CS, and GenEpi. Five methods, except GenEpi, are selected to produce PRS prediction from the external summary statistics without individual genetic data. On the other hand, GenEpi method is included due to its consideration of gene-based epistasis, which is a distinct machine learning-based algorithm to estimate PRS, for comparison.

#### Clumping and Thresholding

Clumping and thresholding (C+T) is the classical algorithm that adjusts the LD using clumping and selects SNPs with *P*-value less than a specified threshold to calculate the PRS for each individual^10^. In PGSbuilder, SNPs within 250 kb away from the index SNP and have the R-squared over 0.1 with it are assigned to the clump of the index SNP. Nine thresholds, including 10^−8^, 10^−7^, 10^−6^, 10^−5^, 10^−4^, 10^−3^, 10^−2^, 10^−1^, and 1, are applied to the clumped SNPs to build PRS models. Beta scores derived from the summary statistics are set as the effect size estimates directly. The model with the best performance on the target set is selected as the final PRS model.

#### PRSice2

PRSice2 is also a clumping and thresholding-based PRS algorithm with a higher resolution of thresholds^17^. SNPs with a minor allele frequency lower than 0.01 are filtered out. Like C+T, beta scores are set as the effect size estimates directly.

#### Lassosum

Lassosum uses penalized regression to adjust the effect size of SNPs for a PRS model^16^. The summary statistics provide the SNP-wise correlation with the phenotype and the initial effect size of SNPs. LD blocks are defined from the subpopulation of the 1000 Genome database, and the LD matrix is calculated from the target set. Additionally, the target set is used for the selection of hyperparameters to get the best PRS model.

#### LDpred2

LDpred2 is a Bayesian PRS predictor by adjusting the effect size of SNPs from the summary statistics^12^. The target set provides the correlation between SNPs for LD estimation within 3 centimorgan. In PGSbuilder, for summary statistics having more than 10 SNPs with *P*-value<10^−8^, we implement the “LDpred2-grid” mode to select the best hyperparameters, including the proportion of causal variants and the heritability. On the other hand, for those with less significant SNPs, we implement the “LDpred2-inf” mode, an infinitesimal model.

#### PRS-CS

PRS-CS is a Bayesian polygenic prediction method that infers the posterior effect size of SNPs from the summary statistics using continuous shrinkage priors^13^. In PGSbuilder, we use the 1000 Genome dataset as the reference panel for LD estimation. The global shrinkage parameter is fixed at 0.2 and other parameters are left as defaults.

#### GenEpi

GenEpi, a machine learning approach, takes both additive effect and SNP-SNP interactions into consideration to build a PRS model from the raw genomic data^36^. GenEpi uses two-stage feature selection to select a single SNP, intragenic interaction, and intergenic interaction and then applies a regression model to fit the selected features. In PGSbuilder, we only train the GenEpi model on the base set.

### Covariates

In GWAS analysis, covariates are used to adjust the genetic effect on the target phenotype. PGSbuilder performs PCA before GWAS, and the top ten principal components (PCs) are served as covariates. In addition, users can provide a covariate file, and covariates with the variance inflation factor (VIF) less than 50 or a missing rate over 20% are removed. Finally, the effect size of each SNP is corrected with PCs and provided covariates during the association test.

On the other hand, to provide a comprehensive risk assessment for individuals, features other than genetic factors should be taken into consideration. After building a PRS model, PGSbuilder combines the PRS score as a genetic factor and user-provided covariates as clinical factors to build a regression model trained on the target set. Then, PGSbuilder predicts each individual using this regression model to stratify the risk of the target phenotype.

### Variant annotation tools

The annotation of significant SNP from GWAS or other genomic analysis is of great importance. Annotation of variants is vital for the translation of genomic results to the functional level for further analysis. The Ensembl Variant Effect Predictor (VEP) is an open-source, powerful, and versatile toolset for the annotation and prioritization of genomic variants for a transcript or even non-coding region^28^. We select VEP (version 106) because of its broad collection of databases, scalability, and free open license. In order to display the important variant information to show first on the web page, PGSbuilder sorts the VEP results by several criteria, including transcript consensus, mutation consequence, mutation severity, and feature biotype. The complete VEP result is provided in the downloaded file. In addition, allele frequencies from Taiwan Biobank^37^ and 1000 Genome Project are provided in the VEP annotation.

Moreover, we integrate our literature mining engines, variant2literature^38^ and pubmedKB^29^, by retrieving entity mentions and odds ratio statistics to create a report of textual evidence for each variant-phenotype pair. The literature report contains an overall summary and single paper snippets. For the overall summary, we first collect sentences and clinical case sentences where the target variant and phenotype are both mentioned. We then present the most important sentences and clinical cases identified by page rank^39^. For single paper snippets, we present the paragraph describing odds ratio statistics of the target variant and phenotype.

### Example Data

#### Taiwan Biobank

Taiwan Biobank (TWB) is a prospective cohort study with genomic data and a variety of phenotypes collected from Taiwanese population^37^. The TWB cohort contains 27,500 individuals genotyped for 653,288 SNPs on the TWB v1.0 array as well as 68,978 individuals genotyped for 748,344 SNPs on the TWB v2.0 array.

#### NIA ADC Cohort

The NIA ADC Cohort consists of individuals evaluated clinically from National Institute on Aging (NIA)-funded Alzheimer Disease Centers (ADC)^40^. Inclusion criteria of late-onset Alzheimer’s disease are autopsied subjects with age >60 or cases diagnosed with DSM-IV or Clinical Dementia Rating >1^40^. All the seven ADC datasets downloaded from NIAGADS (https://www.niagads.org/datasets) were merged directly as a joint analysis. In total, there are 10,256 samples, including 5,334 cases, 3,973 controls, and 949 unknowns, genotyped for 914,402 SNPs.

## Results

### PGSbuilder analysis workflow

PGSbuilder is a web-based server to provide end-to-end analysis for genetic cohort data including GWAS, PRS, and variant annotation. The GWAS analysis aims to figure out the significant SNPs associated with a specific phenotype while the PRS analysis aims to build a model for the estimation of the individual risk. After the analysis, SNP-level annotation and literature exploration using pubmedKB^29^ are performed to provide useful insights into causal variants.

The GWAS pipeline (Figure 1), which is applied to the whole input dataset, consists of quality control (QC) and association tests. As for the PRS pipeline (Figure 1), the input dataset is firstly separated into training and test sets. The training set is undergone QC steps and split into base and target subsets. The base subset is used to obtain the summary statistics of GWAS, while the target subset is used to build the PRS models. On the other hand, the option of an external summary statistics file is available in PGSbuilder. When the external summary statistics file is provided, it replaces the base subset to provide GWAS results and the entire training set serves as the target subset alternatively. To build a PRS model, most PRS methods combine the summary statistics providing the initial SNP effect sizes with the linkage disequilibrium (LD) estimation derived from the target subset. Of note, GenEpi is unavailable for building a PRS model from the external summary statistics. Finally, to validate the model performance, the estimated risks of individuals in the test set are independently calculated by the adjusted effect size.

**Figure 1.**
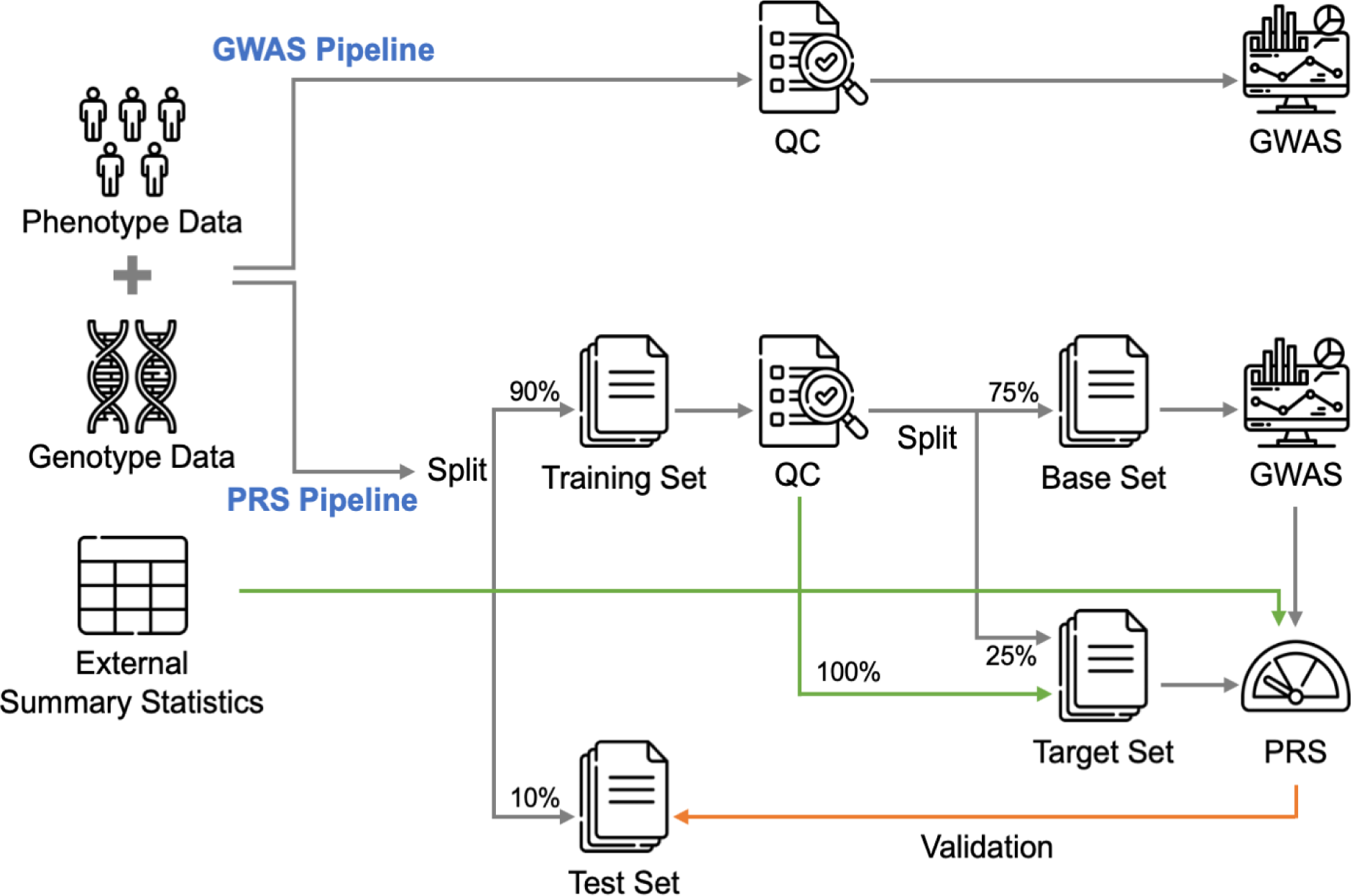
Analysis pipelines of PGSbuilder. PGSbuilder performs GWAS and PRS analysi respectively on the input dataset. For the GWAS pipeline, PGSbuilder applies QC followed by GWAS on the whole input dataset. For the PRS pipeline, PGSbuilder splits the input dataset into training and test sets with the default ratio of 9:1 and applies QC on the training set. The training set is later split into base and target subsets with a ratio of 3:1, and the GWAS result is obtained from the base set. Combining the target set with the summary statistics derived from the base set, PGSbuilder builds PRS models based on different PRS methods. Alternatively, users could provide external summary statistics and the entire training set will be used to build the PRS model. Finally, the independent test set is used to evaluate the performance of the PRS model.

### System implementation

We used Kubernetes and docker technology to group our applications including web interfaces, data processing, GWAS and PRS pipelines, and variant annotation into a service platform. For the web interface, we adopted React architecture and Node.js for the frontend and backend respectively. For the analysis, after users upload genotype data, PGSbuilder will create pods for GWAS and PRS pipelines dynamically and instantly. Significant variants derived from GWAS and PRS pipelines will be annotated through the VEP and pubmedKB to determine the effect of variants in the public database and academic literature.

For the security of private genomic data, users have to sign up via email activation. After login, two studies including a binary trait (classification model) and a quantitative trait (regression model) are demonstrated on the analysis page. To create a new study, users have to upload genotype data in PLINK format and fill in relevant information such as population, genome build, and prediction method (classification or regression). PGSbuilder provides flexibility for users to modify some quality control parameters and select multiple PRS methods (Figure 2A). If the data is successfully uploaded to the PGSbuilder server, the job is added to the analysis queue and will be processed as soon as possible. Users will receive an email notice to check the state of jobs on the running page. Once the job is completed, users can download a comprehensive report for GWAS and PRS results. PGSbuilder also provides an interactive interface to view the result in detail.

**Figure 2.**
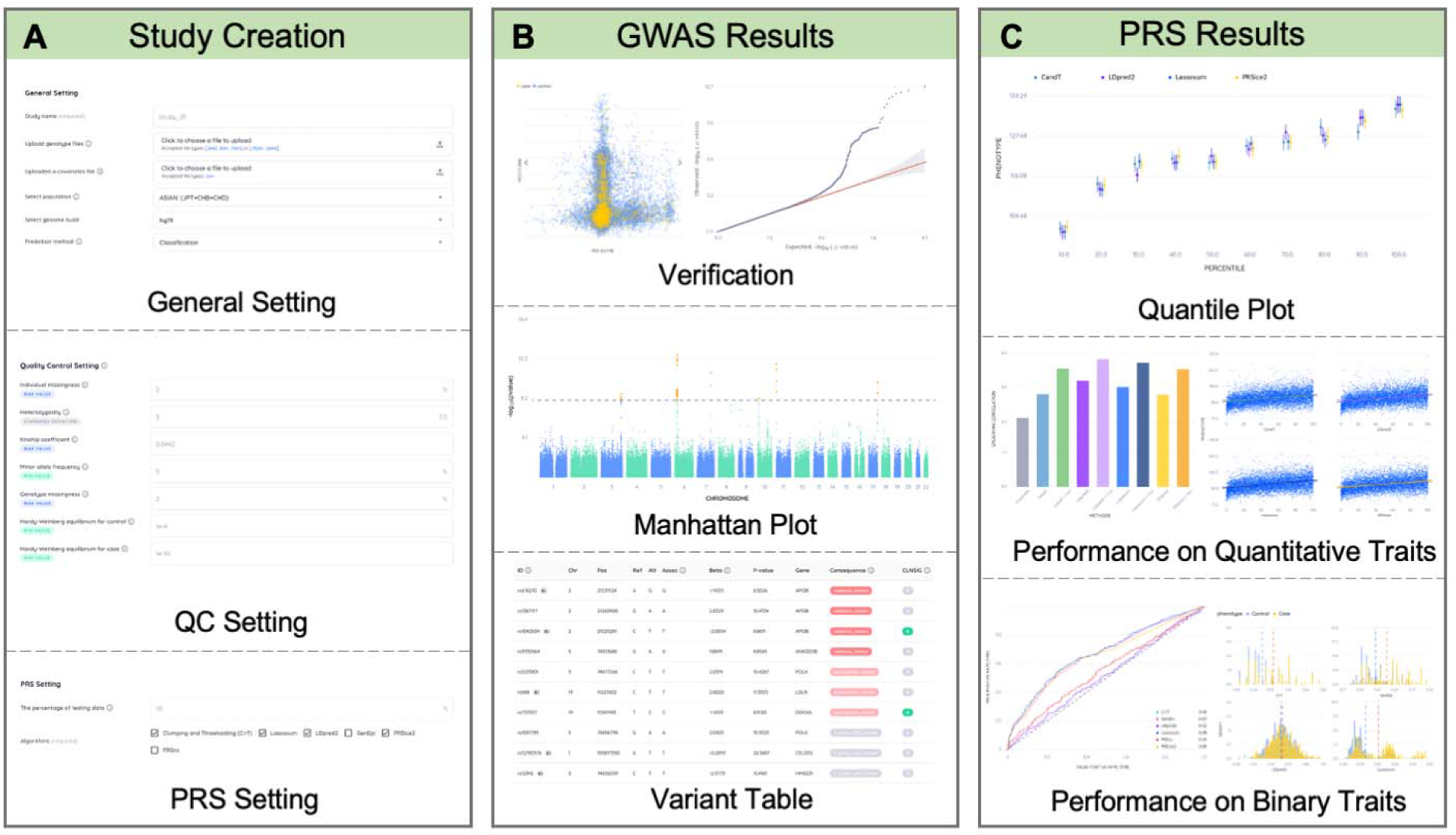
PGSbuilder interface and visualizations. (A) First of all, users can create a new study with customization, including general, QC, and PRS settings. (B) After analysis, GWAS results are composed of verification, including the PCA and Q-Q plots, and significant SNPs, including the Manhattan plot and variant table. (C) On the other hand, PRS results show a quantile plot for risk stratification and performance comparison for quantitative or binary traits.

On the GWAS result page, PCA plot, quantile-quantile plot (Q-Q plot), Manhattan plot, and the variant table are demonstrated (Figure 2B). PCA is used for the correction of population stratification, and the top 10 principal components (PCs) are selected as covariates for GWAS. The paired distributions of the top 3 PCs are shown interactively, and users can arbitrarily switch between three figures through arrow buttons. In addition, each dot represents a sample whose ID will be displayed via a mouseover event, which can help users discriminate outliers. The Q-Q plot is provided to evaluate the deviation of observed *P*-values from expected *P*-values under a uniform distribution. For the Manhattan plot and variant table, we set a suggestive *P*-value threshold of 1×10^−5^ and a strict *P*-value threshold of 5×10^−8^. SNPs with a *P*-value smaller than the threshold are colored in orange and listed in the variant table. The SNPs in the Manhattan plot and the variant table are interactive. Clicking on an orange point on the Manhattan plot navigates the variant table to the corresponding SNP with its information, and vice versa. Besides, users can search for a specific SNP through the search bar. More detailed information of all SNPs including their *P*-values and annotated information are compressed as a zip file to be downloaded.

On the PRS result page, we compare the performance of selected PRS methods. The quantile plot shows the risk stratification (Figure 2C). For each method, samples in the test set are divided into 10 quantiles of increasing PRS. Then, in each quantile, the odds ratio is calculated for binary phenotypes while the mean of values is calculated for quantitative phenotypes. A great difference between the first and the last group represents a good risk stratification. Of note, all individuals in the test set serve as the baseline for odds ratio calculation for binary tracts. In the classification analysis for a binary tract, the receiver operating characteristic (ROC) curve and distribution plot for each method are demonstrated (Figure 2C). The area under the ROC curve illustrates the performance and the distribution plots illustrate the prediction distribution for cases against controls. In the regression analysis for a quantitative tract, Spearman correlations and scatter plots are shown (Figure 2C). The Spearman correlation is performed to evaluate the performance and the scatter plot with a regression line illustrates the relationship between phenotypes and prediction rankings for each method. The tabs of method lists allow users to switch results between different methods. Users can click one of them to view the corresponding performance and variant table.

Furthermore, analysis beyond genetic factors is also available in PGSbuilder. If the covariate file is provided, covariates will be used to correct the effect size of SNPs during GWAS, and then serve as clinical factors combined with PRSs to build a regression model for risk prediction. The performance with or without clinical factors is also demonstrated in the figures for comparison. The weight of each clinical factor is shown in a table for users to figure out important factors.

### Variant annotation panel

In order to help interpret GWAS and PRS results, PGSbuilder provides a comprehensive variant annotation panel for users to explore biological significance. There are often a large number of SNPs associated with a phenotype. PGSbuilder will automatically sort the important SNPs at the top of the panel according to several annotation information including transcript consensus, mutation consequence, mutation severity, and feature biotype. Figure 3 displays an example of the significant SNP information from the GWAS results. Accordingly, three key features are present including variant effect prediction information, external links about the variant, and the related literature. PGSbuilder uses ClinVar^41^ and VEP^28^ for variant interpretation (Fig. 3B). Several external links are provided to easily navigate the further variant information (Fig. 3C). Lastly, PGSbuilder integrates the literature mining results from the pubmedKB^29^ to assist researchers and clinical professionals in obtaining the related literature.

**Figure 3.**
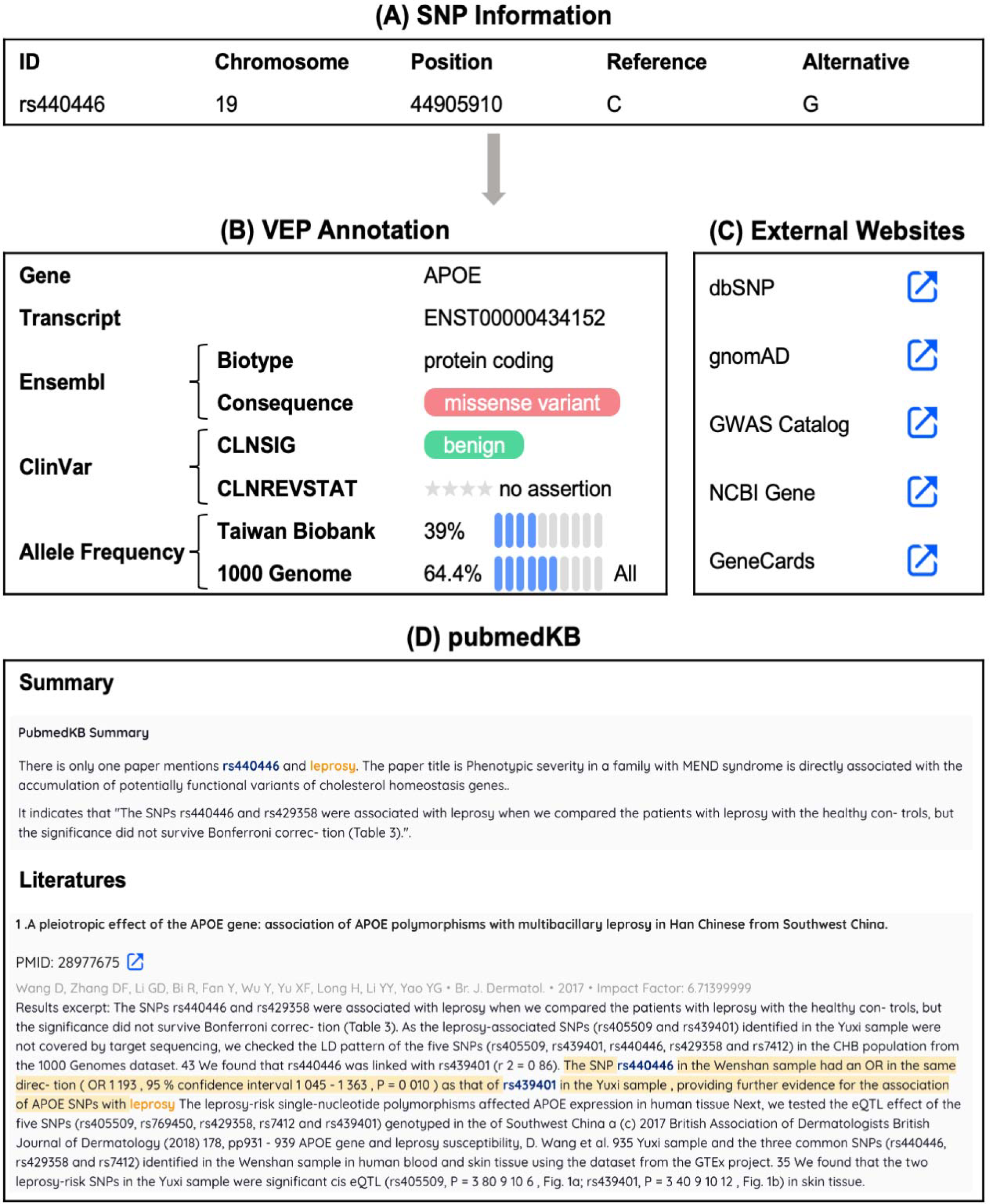
Example annotation result of SNP “rs440446” on PGSbuilder. (A) There is the basic information and statistics (e.g. GWAS *P*-value) of the variant. (B) We apply different colors on consequence (the red one) and ClinVar significance (the green one) according to table provided by Ensembl and ClinVar, respectively,for a better presentation of SNP importance level. The following block is the transcript ID, ClinVar significance, and allele frequency from VEP. (C) We also provided links to external websites with more variant or gene information, such as dbSNP^59^, gnomAD^60^, GWAS Catalog^2^, and GeneCards^61^. (D) The block at the bottom is the results from pubmedKB. The summary presents the sentence where the SNP and the phenotype co-occur, and we show the paper snippet of odds ratio statistics.

### System performance

For benchmarking, we recorded execution time, average memory, and CPU usage for QC, GWAS, and PRS methods with 680k SNPs given 20k, 50k, and 110k samples (Table 1). The resource for each execution was limited to 20 GB and 10 CPUs. Obviously, more resources were needed as the sample size increased. Table 1 shows the comparison between six PRS methods. PRSice2, PRS-CS, and GenEpi took much more execution time than the others, but PRSice2 and GenEpi used the least CPU and memory respectively. In conclusion, it takes about three days to complete a comprehensive PRS analysis for a dataset with 110k samples and 680k SNP.

**Table 1.** The system performance, including execution time, average CPU, and memory of PGSbuilder. We performed QC, GWAS, and six PRS methods (classification for a binary trait) on a dataset with the same number of SNPs but different sample sizes.

| Sample | STATS | QC | GWAS | C+T | Lassosum | LDpred2 | PRSice2 | PRS-CS | GenEpi | Total |
| --- | --- | --- | --- | --- | --- | --- | --- | --- | --- | --- |
| 20k | Time (min) | 8.0 | 8.0 | 15.5 | 18.8 | 27.3 | 79.2 | 171.8 | 450.9 | 779.5 |
| 20k | Avg. CPU | 2.9 | 6.9 | 5.6 | 4.7 | 6.3 | 3.3 | 7.5 | 7.7 |  |
| 20k | Memory (GB) | 5.1 | 2.0 | 10.9 | 10.5 | 10.9 | 10.7 | 8.4 | 3.8 |  |
| 50k | Time (min) | 43.5 | 20.0 | 33.4 | 41.6 | 73.9 | 180.8 | 268.6 | 715.0 | 1376.8 |
| 50k | Avg. CPU | 4.7 | 6.8 | 7.9 | 7.6 | 7.6 | 5.6 | 8.0 | 7.4 |  |
| 50k | Memory (GB) | 16.0 | 1.7 | 16.2 | 16.1 | 15.8 | 14.8 | 12.9 | 3.8 |  |
| 110k | Time (min) | 139.5 | 77.0 | 200.3 | 220.3 | 251.3 | 510.6 | 967.3 | 2086.9 | 4453.1 |
| 110k | Avg. CPU | 4.8 | 8.6 | 7.4 | 7.3 | 7.6 | 5.5 | 8.0 | 7.2 |  |
| 110k | Memory (GB) | 19.4 | 13.8 | 17.0 | 17.0 | 16.7 | 13.9 | 12.0 | 10.6 |  |

### Case Study

To demonstrate the capability of PGSbuilder, we performed two case studies using the cohorts with a large number of individuals and corresponding phenotypes. Firstly, in the Taiwan Biobank (TWB)^37^, a Taiwanese cohort composed of healthy adults, we previously defined nine quantitative traits and five binary traits related to some common chronic diseases, such as type 2 diabetes or dyslipidemia, according to their phenotypic measures (see https://github.com/chienyuchen/TWB-PRS for more information). The presented GWAS and PRS models across fourteen traits in the TWB were built by using PGSbuilder. Among them, low-density lipoprotein (LDL), a quantitative trait, was selected here to demonstrate the usage of adding covariates and the leverage of external summary statistics to run PGSbuilder. Secondly, for the cohort with a specific disease, we performed GWAS and PRS analysis on the National Institute on Aging (NIA)-funded Alzheimer Disease Centers (ADC) Cohort^40^ to demonstrate the result of a binary trait.

#### Low-density lipoprotein

Low-density lipoprotein (LDL), which is a kind of lipoprotein to transport fat molecules around the body, acts as the primary driver of atherogenesis resulting in cardiovascular diseases^42^. Several genes, such as LDLR, PCSK9, and APOB, affecting the quantity of LDL in circulation have been reported^43^. Recognizing people with a genetic tendency for high LDL could help them by providing early intervention to avoid the progression of severe cardiovascular diseases. Therefore, in this study, we applied GWAS and PRS analysis using PGSbuilder on the TWB data. The covariates, including age, sex, and body mass index (BMI), were added to correct GWAS for genetic factors and then serve as clinical factors to build regression models for risk prediction.

With the default QC settings of PGSbuilder, 55,412 samples and 276,068 SNPs were passed the quality control (Table S1-2). To control the population stratification, PGSbuilder always performs PCA analysis and applies the top ten principal components (PCs) as covariates during GWAS. Figure 4A demonstrates the distribution of PC1 and PC2 to confirm SNPs without unusual differentiation between quantiles in the TWB data. The interactive Manhattan plot is shown in Figure 4B and the significant SNPs with a *P*-value < 10^−5^ are highlighted in orange for clicking to navigate variant information. Notably, in comparison with the previous study using the same TWB data^44^, highly similar results were observed in PGSbuilder as shown that more than 80% (89/111) of significant SNPs in the TWB arrays were identically found to associate with the LDL trait. That is, the pipeline in PGSbuilder is indeed reproducible.

**Figure 4.**
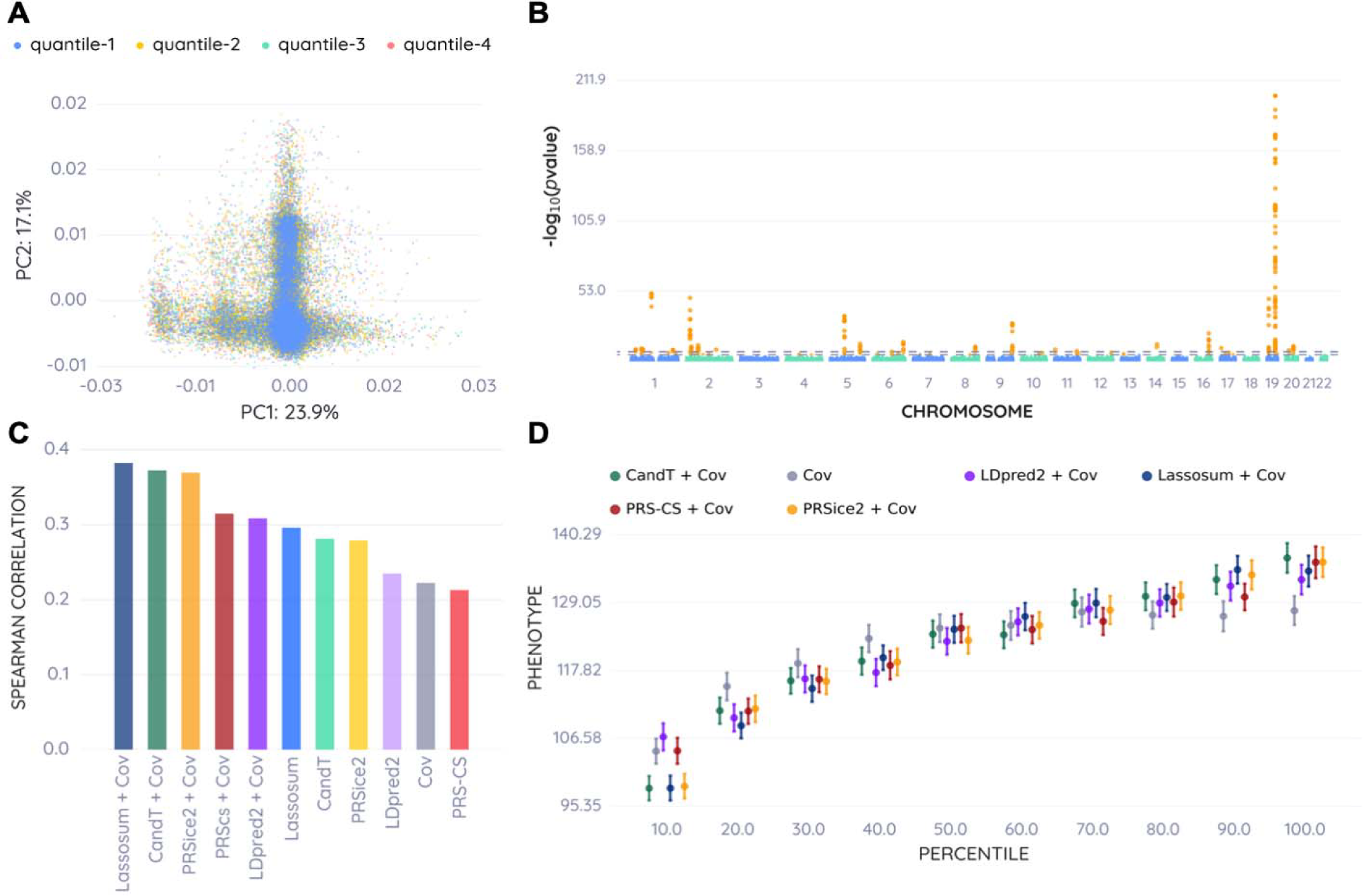
Results of LDL GWAS and PRS analyses. (A) PCA plot of the first and second PCs. To view any deviation of PCs among the samples, values of the quantitative phenotype are separated into four quantiles. (B) Manhattan plot of −log10(P-value). GWAS is performed on autosomal SNPs, and SNPs with P-value <10-5 are colored in orange. (Source data in Table S3) (C) Bar plot of Spearman’s correlation of each PRS model. Models derived from different methods with or without covariates (Cov) are demonstrated simultaneously. (Source data in Table S4) (D) Quantile plot for risk stratification. The “covariate-only (Cov)” model and “PRS + covariate” models are plotted to compare the usage of genetic factors. (Source data in Tabl S5).

In addition, PGSbuilder allows users to provide external summary statistics to build PRS models. Herein, the external summary statistics from the BioBank Japan^45^ to identify significant variants and stratify people by the risk of high LDL were applied to estimate PRS in the TWB data. Figure 4C shows the performance on the test set of each PRS method with and without clinical factors. Overall, PRS combined with clinical factors performs better than PRS-only and clinical factors-only models. These results indicate that the genetic factor combined with clinical factors provide a better prediction effect. Figure 4D depicts the risk stratification of models using clinical factors. “PRS + clinical factors” models stratified the test set better than the “clinical factors-only” model. In the “PRS + clinical factors” models, the difference in average LDL between the first and last groups is up to forty. Furthermore, the weight of each feature in the “PRS-clinical factors” model is listed in Table 2, where PRS has the largest contribution in all the models.

**Table 2.**
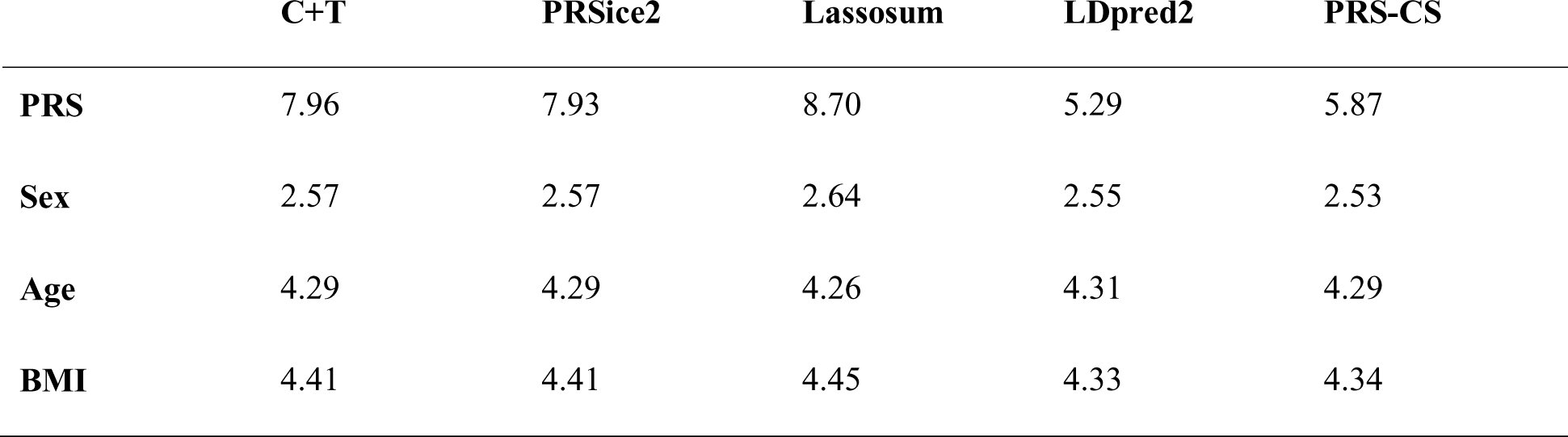
The weight of PRS and clinical factors for “PRS + clinical factors” models of LDL.<colcnt=6>

#### Alzheimer’s disease

Alzheimer’s disease (AD), the major cause of dementia, is a complex disorder associated with genetic factors and environmental factors^46^. Several genetic loci, such as APOE, have been identified at the level of association study^47, 48^. Combining the effects of these genetic loci to build a PRS model could provide individuals with the disease risk for further preventive strategies^49^. In this study, to build PRS models based on different methods and compare the performance of them, we analyzed the National Institute on Aging (NIA)-funded Alzheimer Disease Centers (ADC) cohort using PGSbuilder.

Figure 5 shows the performance of PRS analysis from PGSbuilder. There are two obvious groups with different performances. C+T, PRSice2, Lassosum, and GenEpi have better auROC than LDpred2 and PRS-CS (Figure 5A). Figure 5B depicts the prediction distribution of cases and controls; the more distance between the distributions the better performance of the model. For further comparison of different methods, an UpSet plot depicts the intersection of top-100 valuable SNPs from each method (Figure 5C). Notably, LDpred2 and PRS-CS have some distinct SNPs than others, which might cause noise for the PRS prediction and decrease the model performance.

**Figure 5.**
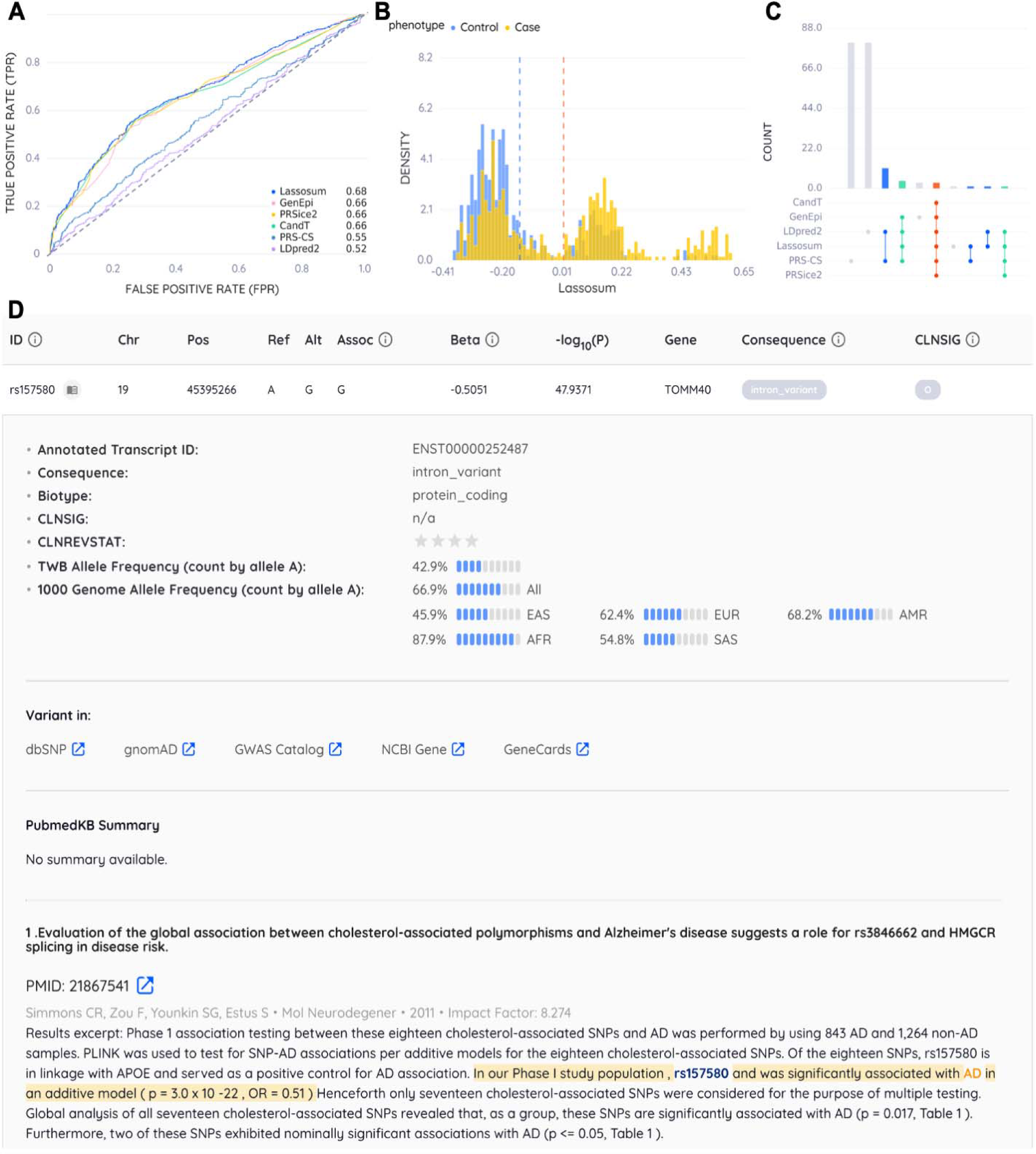
R**esults of AD across different PRS methods. (**A) ROC curve of each PRS model on the test set. (Source data in Table S6) (B) Prediction distributions of the Lassosum PRS model for cases (yellow) and controls (blue). The dashed line represents the mean of each group. (C) UpSet plot^62^ for the intersection of important SNPs derived from different PRS methods. The intersection, or the combination, of methods are presented as the matrix layout while the variant counts of each intersection are shown as the histogram. Different colors represent the number of PRS methods. (corresponding output data in Table S7) (D) Annotations for SNP “rs157580”. On the top is the basic information and statistics of the variant. The following block is the transcript ID, ClinVar significance and allele frequency from VEP^28^. In addition, we also provided links to external websites with more variant information, such as dbSNP^59^ and gnomAD^60^. The block in the bottom is the results from pubmedKB^29^ which highlights the odds ratio of AD in the presence of this variant.

To investigate the information of SNPs, PGSbuilder annotates SNPs using VEP^28^ and pubmedKB^29^. For example, Figure 5D shows the annotation of rs157580, which is an intron variant of gene TOMM40 with average allele frequency across different populations. A previous study (PMID: 21867541) also reported that rs157580 was significantly associated with AD^50^. The literature mining of PubMed abstracts by pubmedKB facilitates users to interpret the variants more readily.

## Discussion

PGSbuilder is a cloud-based platform that offers comprehensive genotyping analyses, including GWAS and PRS, all in one place. Our goal for GWAS is to help identify significant SNPs associated with the target phenotype, while for PRS, we aim to assist evaluation of the prediction performance of polygenic models. Customized settings are available for users to adjust the analytic process, such as quality control, population stratification, and the selection of PRS methods. With PGSbuilder’s interactive interfaces, users can easily interpret their results. For instance, users can select specific SNPs on the Manhattan plot and view the corresponding annotations in the table. Additionally, PGSbuilder integrates pubmedKB for variant interpretation by providing literature support. With these features, PGSbuilder is a comprehensive and user-friendly platform for GWAS and PRS.

In addition to the analytic pipeline, PGSbuilder offers various visualization plots to compare the performance of different PRS methods. To evaluate risk stratification, the quantile plot is a key interpretation tool. The UpSet plot enables users to observe the intersection of important SNPs selected from each method. Additionally, PGSbuilder incorporates our original GenEpi software^36^, which provides a unique method to uncover the genetic epistasis associated with phenotypes, as demonstrated in other recent studies^51, 52^. Finally, as clinical factors are provided, PGSbuilder will rank the weights of them and PRS to highlight the most predictive feature, which helps users investigate the risk factor precisely.

While PGSbuilder provides a range of useful features, there are some limitations to its functionality. First, it is important to consider the limitations of hardware resources when dealing with large datasets. For example, some imputed files containing 10 million SNPs and 50K samples may not be immediately accessible due to these restrictions. However, computationally efficient methods such as C+T, Lassosum, and PRSice2 can eb effectively applied to such datasets, based on our internal experiments. It is worth noting that building a predictive model using some PRS methods may require a significant amount of time. On the other hand, GenEpi, which discovers the gene-based epistasis, is not practical for imputed data due to its computational complexity. Secondly, some known PRS methods, such as those based on a mixture model for SNP effective size (e.g. SBayesR^14^, DPR^53^, DBSLMM^54^), are currently not included in PGSbuilder. Lastly, PRS models can only be downloaded from PGSbuilder output directly. Going forward, we are planning to implement a prediction module that allows users to upload other datasets and then automatically obtain predictions of available PRS models .

The field of PRS development is growing rapidly, with mounting evidence using the wealth of data collected in biobanks^55–58^. As the proof of concept is solidly demonstrated, an effective and comprehensive platform is necessary to perform GWAS and PRS analysis for diseases that are not covered by biobanks. PGSbuilder provides researchers with the ability to identify significant loci with annotations and investigate the polygenicity of a target phenotype across a specific population effectively. By leveraging genotypes, a PRS model has the clinical potential to offer risk evaluations to individuals. This, in turn, can facilitate early surveillance for severe diseases.

## Conclusion

PGSbuilder is an end-to-end platform that seamlessly integrates QC of genotype data, GWAS, PRS, SNP annotation, and visualizations. This platform is versatile, allowing the incorporation of external GWAS summary statistics to run PRS using various methods, thereby enabling the estimation of genetic risk in smaller cohort samples. In addition, PGSbuilder’s user-friendly interface is designed to be accessible to users without programming experiences. In the future, we plan to further augment and broaden PGSbuilder by introducing a prediction module that allows users to directly run their PRS models for specific disease phenotypes.

## Acknowledgements

We thank Tzu-Hung Hsiao and Chien-Lin Mao for valuable early discussion and pipeline testing.

## Data and software availability

All genetic and phenotype data in TWB described in this paper are publicly available via the Taiwan Biobank data access protocol. Fourteen PRS models using TWB data, including five binary phenotypes and nine quantitative traits, are freely available on the GitHub project repository (https://github.com/chienyuchen/TWB-PRS). The AD data is publicly available to registered researchers by request from the National Institute on Aging Genetics of Alzheimer’s Disease Data Storage Site (NIAGADS). The source codes for GWAS and PRS analyses were deposited to Github and is available at https://github.com/ailabstw/PGSbuilder.

## Ethics approval and consent to participate

The application number of TWB data is TWBR10411-03. This application of NIA ADC Cohort dataset has been filed with the IRB (202106049RINA) in order to get approval from NIAGADS.

## Authors’ contributions

KHL, YLL, TTH, YCC, and HCC conceived and implemented the pipeline development. YCC inspired team members to unite as a product manager, and designed all the frameworks of this web service, including wireframe, prototype, and database schema. SSW, WCL, and GZF implemented the web design and interface. TFC and PHL implemented the literature mining. YLK served as liaisons to user communities. YCC and JHH helped project development and management. PLC led the application of TWB data. HFJ, HKT, CYC, and JHH supervised the project. KHL and JHH led the writing of the manuscript. All authors discussed the results and implications and commented on the manuscript. All authors read and approved the final manuscript.

## References

1. Mills, M. C. & Rahal, C. A scientometric review of genome-wide association studies. Commun Biol 2, 9 (2019).

2. Buniello, A. et al. The NHGRI-EBI GWAS Catalog of published genome-wide association studies, targeted arrays and summary statistics 2019. Nucleic Acids Research vol. 47 D1005–D1012 Preprint at https://doi.org/10.1093/nar/gky1120 (2019).

3. Visscher, P. M. et al. 10 Years of GWAS Discovery: Biology, Function, and Translation. Am. J. Hum. Genet. 101, 5–22 (2017).

4. Stranger, B. E., Stahl, E. A. & Raj, T. Progress and promise of genome-wide association studies for human complex trait genetics. Genetics 187, 367–383 (2011).

5. Watanabe, K. et al. A global overview of pleiotropy and genetic architecture in complex traits. Nat. Genet. 51, 1339–1348 (2019).

6. Chatterjee, N., Shi, J. & García-Closas, M. Developing and evaluating polygenic risk prediction models for stratified disease prevention. Nat. Rev. Genet. 17, 392–406 (2016).

7. Wray, N. R. et al. From Basic Science to Clinical Application of Polygenic Risk Scores: A Primer. JAMA Psychiatry 78, 101–109 (2021).

8. Ma, Y. & Zhou, X. Genetic prediction of complex traits with polygenic scores: a statistical review. Trends Genet. 37, 995–1011 (2021).

9. Wray, N. R., Goddard, M. E. & Visscher, P. M. Prediction of individual genetic risk to disease from genome-wide association studies. Genome Res. 17, 1520–1528 (2007).

10. International Schizophrenia Consortium et al. Common polygenic variation contributes to risk of schizophrenia and bipolar disorder. Nature 460, 748–752 (2009).

11. Vilhjálmsson, B. J. et al. Modeling Linkage Disequilibrium Increases Accuracy of Polygenic Risk Scores. Am. J. Hum. Genet. 97, 576–592 (2015).

12. Privé, F., Arbel, J. & Vilhjálmsson, B. J. LDpred2: better, faster, stronger. Bioinformatics (2020) doi:10.1093/bioinformatics/btaa1029.

13. Ge, T., Chen, C.-Y., Ni, Y., Feng, Y.-C. A. & Smoller, J. W. Polygenic prediction via Bayesian regression and continuous shrinkage priors. Nat. Commun. 10, 1776 (2019).

14. Lloyd-Jones, L. R. et al. Improved polygenic prediction by Bayesian multiple regression on summary statistics. Nat. Commun. 10, 5086 (2019).

15. Zhou, G. & Zhao, H. A fast and robust Bayesian nonparametric method for prediction of complex traits using summary statistics. PLoS Genet. 17, e1009697 (2021).

16. Mak, T. S. H., Porsch, R. M., Choi, S. W., Zhou, X. & Sham, P. C. Polygenic scores via penalized regression on summary statistics. Genet. Epidemiol. 41, 469–480 (2017).

17. Choi, S. W. & O’Reilly, P. F. PRSice-2: Polygenic Risk Score software for biobank-scale data. Gigascience 8, (2019).

18. Ni, G. et al. A Comparison of Ten Polygenic Score Methods for Psychiatric Disorders Applied Across Multiple Cohorts. Biol. Psychiatry 90, 611–620 (2021).

19. Pain, O. et al. Evaluation of polygenic prediction methodology within a reference-standardized framework. PLoS Genet. 17, e1009021 (2021).

20. Collister, J. A., Liu, X. & Clifton, L. Calculating Polygenic Risk Scores (PRS) in UK Biobank: A Practical Guide for Epidemiologists. Front. Genet. 13, 818574 (2022).

21. Choi, S. W., Mak, T. S.-H. & O’Reilly, P. F. Tutorial: a guide to performing polygenic risk score analyses. Nat. Protoc. 15, 2759–2772 (2020).

22. Wray, N. R. et al. Research review: Polygenic methods and their application to psychiatric traits. J. Child Psychol. Psychiatry 55, 1068–1087 (2014).

23. Lambert, S. A. et al. The Polygenic Score Catalog as an open database for reproducibility and systematic evaluation. Nat. Genet. 53, 420–425 (2021).

24. Martin, A. R. et al. Clinical use of current polygenic risk scores may exacerbate health disparities. Nat. Genet. 51, 584–591 (2019).

25. Scutari, M., Mackay, I. & Balding, D. Using Genetic Distance to Infer the Accuracy of Genomic Prediction. PLoS Genet. 12, e1006288 (2016).

26. Wang, Y. et al. Theoretical and empirical quantification of the accuracy of polygenic scores in ancestry divergent populations. Nat. Commun. 11, 3865 (2020).

27. Folkersen, L. et al. Impute.me: An Open-Source, Non-profit Tool for Using Data From Direct-to-Consumer Genetic Testing to Calculate and Interpret Polygenic Risk Scores. Front. Genet. 11, 578 (2020).

28. McLaren, W. et al. The Ensembl Variant Effect Predictor. Genome Biol. 17, 122 (2016).

29. Li, P.-H. et al. pubmedKB: an interactive web server for exploring biomedical entity relations in the biomedical literature. Nucleic Acids Res. (2022) doi:10.1093/nar/gkac310.

30. Chang, C. C. et al. Second-generation PLINK: rising to the challenge of larger and richer datasets. Gigascience 4, 7 (2015).

31. Manichaikul, A. et al. Robust relationship inference in genome-wide association studies. Bioinformatics 26, 2867–2873 (2010).

32. Consortium, T. I. H. 3. & The International HapMap 3 Consortium. Integrating common and rare genetic variation in diverse human populations. Nature vol. 467 52–58 Preprint at https://doi.org/10.1038/nature09298 (2010).

33. Marees, A. T. et al. A tutorial on conducting genome-wide association studies: Quality control and statistical analysis. International Journal of Methods in Psychiatric Research 27, e1608 (2018).

34. Price, A. L. et al. Principal components analysis corrects for stratification in genome-wide association studies. Nat. Genet. 38, 904–909 (2006).

35. Purcell, S. et al. PLINK: a tool set for whole-genome association and population-based linkage analyses. Am. J. Hum. Genet. 81, 559–575 (2007).

36. Chang, Y.-C. et al. GenEpi: gene-based epistasis discovery using machine learning. BMC Bioinformatics 21, 68 (2020).

37. Feng, Y.-C. A. et al. Taiwan Biobank: a rich biomedical research database of the Taiwanese population. Preprint at https://doi.org/10.1101/2021.12.21.21268159.

38. Lin, Y.-H. et al. variant2literature: full text literature search for genetic variants. bioRxiv 583450 (2019) doi:10.1101/583450.

39. Page, L., Brin, S., Motwani, R. & Winograd, T. The PageRank Citation Ranking: Bringing Order to the Web. (1999).

40. Naj, A. C. et al. Common variants at MS4A4/MS4A6E, CD2AP, CD33 and EPHA1 are associated with late-onset Alzheimer’s disease. Nat. Genet. 43, 436–441 (2011).

41. Landrum, M. J. et al. ClinVar: improving access to variant interpretations and supporting evidence. Nucleic Acids Res. 46, D1062–D1067 (2018).

42. Borén, J. et al. Low-density lipoproteins cause atherosclerotic cardiovascular disease: pathophysiological, genetic, and therapeutic insights: a consensus statement from the European Atherosclerosis Society Consensus Panel. Eur. Heart J. 41, 2313–2330 (2020).

43. Borén, J. et al. Low-density lipoproteins cause atherosclerotic cardiovascular disease: pathophysiological, genetic, and therapeutic insights: a consensus statement from the European Atherosclerosis Society Consensus Panel. Eur. Heart J. 41, 2313–2330 (2020).

44. Chen, C.-Y. et al. Analysis across Taiwan Biobank, Biobank Japan and UK Biobank identifies hundreds of novel loci for 36 quantitative traits. Preprint at https://doi.org/10.1101/2021.04.12.21255236.

45. Sakaue, S. et al. A cross-population atlas of genetic associations for 220 human phenotypes. Nat. Genet. 53, 1415–1424 (2021).

46. Breijyeh, Z. & Karaman, R. Comprehensive Review on Alzheimer’s Disease: Causes and Treatment. Molecules 25, (2020).

47. Wightman, D. P. et al. A genome-wide association study with 1,126,563 individuals identifies new risk loci for Alzheimer’s disease. Nat. Genet. 53, 1276–1282 (2021).

48. Bellenguez, C. et al. New insights into the genetic etiology of Alzheimer’s disease and related dementias. Nat. Genet. 54, 412–436 (2022).

49. de Rojas, I. et al. Common variants in Alzheimer’s disease and risk stratification by polygenic risk scores. Nat. Commun. 12, 3417 (2021).

50. Simmons, C. R., Zou, F., Younkin, S. G. & Estus, S. Evaluation of the global association between cholesterol-associated polymorphisms and Alzheimer’s disease suggests a role for rs3846662 and HMGCR splicing in disease risk. Mol. Neurodegener. 6, 62 (2011).

51. Yashin, A. I. et al. Roles of interacting stress-related genes in lifespan regulation: insights for translating experimental findings to humans. J Transl Genet Genom 5, 357–379 (2021).

52. Rodrigo, L. M. & Nyholt, D. R. Imputation and Reanalysis of ExomeChip Data Identifies Novel, Conditional and Joint Genetic Effects on Parkinson’s Disease Risk. Genes 12, (2021).

53. Zeng, P. & Zhou, X. Non-parametric genetic prediction of complex traits with latent Dirichlet process regression models. Nat. Commun. 8, 456 (2017).

54. Yang, S. & Zhou, X. Accurate and Scalable Construction of Polygenic Scores in Large Biobank Data Sets. Am. J. Hum. Genet. 106, 679–693 (2020).

55. Zhang, R. et al. Novel disease associations with schizophrenia genetic risk revealed in ∼400,000 UK Biobank participants. Mol. Psychiatry 27, 1448–1454 (2022).

56. Richardson, T. G., Harrison, S., Hemani, G. & Davey Smith, G. An atlas of polygenic risk score associations to highlight putative causal relationships across the human phenome. Elife 8, (2019).

57. Sakaue, S. et al. Trans-biobank analysis with 676,000 individuals elucidates the association of polygenic risk scores of complex traits with human lifespan. Nat. Med. 26, 542–548 (2020).

58. Shen, X. et al. A phenome-wide association and Mendelian Randomisation study of polygenic risk for depression in UK Biobank. Nat. Commun. 11, 2301 (2020).

59. Sherry, S. T. dbSNP: the NCBI database of genetic variation. Nucleic Acids Research vol. 29 308–311 Preprint at https://doi.org/10.1093/nar/29.1.308 (2001).

60. Karczewski, K. J. et al. The mutational constraint spectrum quantified from variation in 141,456 humans. Nature 581, 434–443 (2020).

61. Safran, M. et al. The GeneCards Suite. in Practical Guide to Life Science Databases (eds. Abugessaisa, I. & Kasukawa, T.) 27–56 (Springer Nature Singapore, 2021).

62. Lex, A., Gehlenborg, N., Strobelt, H., Vuillemot, R. & Pfister, H. UpSet: Visualization of Intersecting Sets. IEEE Trans. Vis. Comput. Graph. 20, 1983–1992 (2014).

